# Consistent cerebellar pathway-cognition associations across pre-adolescents & young adults: a diffusion MRI study of 9000+ participants

**DOI:** 10.1101/2025.02.05.636737

**Authors:** Leo R. Zekelman, Suheyla Cetin-Karayumak, Yuqian Chen, Melyssa Almeida, Jon Haitz Legarreta, Jarrett Rushmore, Steve Pieper, Zhou Lan, John E. Desmond, Lissa C. Baird, Nikos Makris, Yogesh Rathi, Fan Zhang, Alexandra J. Golby, Lauren J. O’Donnell

## Abstract

The cerebellum, long implicated in movement, is now recognized as a contributor to higher-order cognition. The cerebellar pathways provide key structural links between the cerebellum and cerebral regions integral to language, memory, and executive function. Here, we present a large-scale, cross-sectional diffusion MRI (dMRI) analysis investigating the relationships between cerebellar pathway microstructure and cognitive performance in over 9,000 participants spanning pre-adolescence (n>8,000 from the ABCD dataset) and young adulthood (n>900 from the HCP-YA dataset). We assessed the microstructure of five cerebellar pathways—the inferior, middle, and superior cerebellar peduncles; the parallel fibers; and input/Purkinje fibers—using three dMRI measures of fractional anisotropy, mean diffusivity, and number of streamlines. Cognitive performance was evaluated using seven NIH Toolbox assessments of language, executive function, and memory. In both datasets, we found numerous significant associations between cerebellar pathway microstructure and cognitive performance. These associations showed a strong correlation across the two datasets (r = 0.47, p < 0.0001), underscoring the reliability of cerebellar dMRI-cognition relationships in pre-adolescents and young adults. In both datasets, the strongest associations were found between the superior cerebellar peduncle and performance on language assessments, suggesting this pathway plays an important role in language function across age groups. In young adults, but not pre-adolescents, parallel fiber microstructure was linked to inhibitory control, suggesting that contributions to attentional processes may emerge or strengthen with maturation. Overall, our findings highlight the important role of cerebellar pathways in cognition and the utility of large-scale datasets for advancing our understanding of brain-cognition relationships.

**Significance:** This study provides strong evidence linking cerebellar pathway tissue microstructure to cognition across large populations of preadolescent children and young adults. By leveraging diffusion MRI tractography and cognitive performance data from two major datasets, we identify significant relationships between cerebellar pathway microstructure and cognitive performance. Importantly, findings are significantly correlated across datasets, pointing to the consistency of these relationships, bridging age groups and acquisitions. These results highlight the cerebellar pathways’ integral role in cognitive functioning and underscore the value of large-scale, population-based studies in advancing our understanding of brain-cognition relationships.

**Classification** 1) Biological Sciences, Neuroscience; 2) Social Sciences, Psychological and Cognitive Sciences

## 1. Introduction

The cerebellum is increasingly recognized for its role in cognition, supported by studies of patients with isolated cerebellar lesions^1^, functional neuroimaging^2^, and cerebellar stimulation experiments^3^. These studies link the cerebellum to diverse cognitive processes including language, memory, attention, and executive function^4^. Connectivity studies show the cerebellum is functionally coupled to the cerebrum, particularly to prefrontal cortices involved in cognition^5^. The cerebellar pathways form multi-synaptic connections between the cerebellum and cerebrum, providing an anatomical basis for their shared role in cognitive functioning^6^. However, our understanding of how these pathways relate to different human cognitive functions remains limited.

The only non-invasive way to map the human cerebellar pathways in vivo is diffusion magnetic resonance imaging (dMRI)^7^. dMRI measures the molecular motion of water in different orientations, enabling the mapping of cerebellar pathways via tractography^8^. The pathways that are frequently studied in dMRI tractography investigations are the cerebellar peduncles that project to and from the cerebellum: the inferior (ICP), middle (MCP), and superior (SCP) cerebellar peduncles^8^. In addition to these pathways, dMRI tractography facilitates the examination of the parallel fibers (PF) that extend across the cerebellar cortex, as well as the input and Purkinje fibers (IP) that contribute to the formation of the medullary white matter of the cerebellum^9–11^. Furthermore, dMRI enables the quantification of tissue microstructure along the cerebellar pathways. Commonly investigated quantitative dMRI measures include fractional anisotropy (FA), mean diffusivity (MD), and the number of streamlines (NoS)^12^; these measures reflect multiple microanatomical components of tissue microstructure (e.g., myelin formation and remodeling, axon diameter and organization)^13^.

Previous studies of the cerebellar pathways have shown that these quantitative dMRI measures of the cerebellar peduncles are associated with cognitive assessments including assessments of language abilities, executive function, learning, memory, attention, and social cognition^14–26^. However, our current understanding of the relationships between dMRI quantitative measures of the cerebellar pathways and performance on cognitive assessments remains limited. Previous studies report inconsistent findings, such as varying associations between reading abilities and cerebellar pathway microstructure in children, attributed to differences in sample sizes, age ranges, and reading assessments^14,15^. In addition, many studies focus on cognitively impaired populations, making it unclear if findings apply to healthy control populations^20,22,23^. Finally, many studies do not account for developmental differences, making it unclear whether findings generalize across childhood and adulthood^16,18^. Studies leveraging larger datasets, non-clinical populations, targeted age ranges, consistent cognitive assessments, and consistent dMRI measures are needed to clarify these relationships. To address these gaps, we examine the relationships between dMRI measures of cerebellar pathways and cognitive performance using two large, non-clinical datasets: the Adolescent Brain and Cognitive Development Study® (ABCD; n = 8,848, 9.9 ± 0.6 years [mean ± SD], 4,242 females) and the Human Connectome Project Young Adult (HCP-YA; n = 945, 28.7 ± 3.7 years [mean ± SD], 518 females)^27–29^. We analyze three dMRI measures (FA, MD, NoS), five cerebellar pathways (ICP, MCP, SCP, IP, PF), and seven cognitive measures describing language, executive function, processing speed, and memory. Harmonized dMRI data, standardized tractography, consistent pathway definitions, and consistent dMRI measures facilitate comparisons and reproducibility across datasets. With a combined sample size exceeding 9,000 participants, this study provides a comprehensive, cross-sectional analysis of cerebellar pathway measures and cognition across pre-adolescent children and adults.

## 2. Results

### 2.1 Consistent identification and quantification of cerebellar pathways

Five cerebellar pathways (ICP, MCP, SCP, PF, and IP) are consistently identified and quantified in both ABCD and HCP-YA datasets using an automated method that parcellates tractography data using an anatomical atlas^11^. This parcellation approach is consistent across the human lifespan and across dataset-specific dMRI acquisitions^11^. **Figure 1A** provides a visualization of the cerebellar pathways in the atlas, an example ABCD participant, and an example HCP-YA participant.

**Figure 1.**
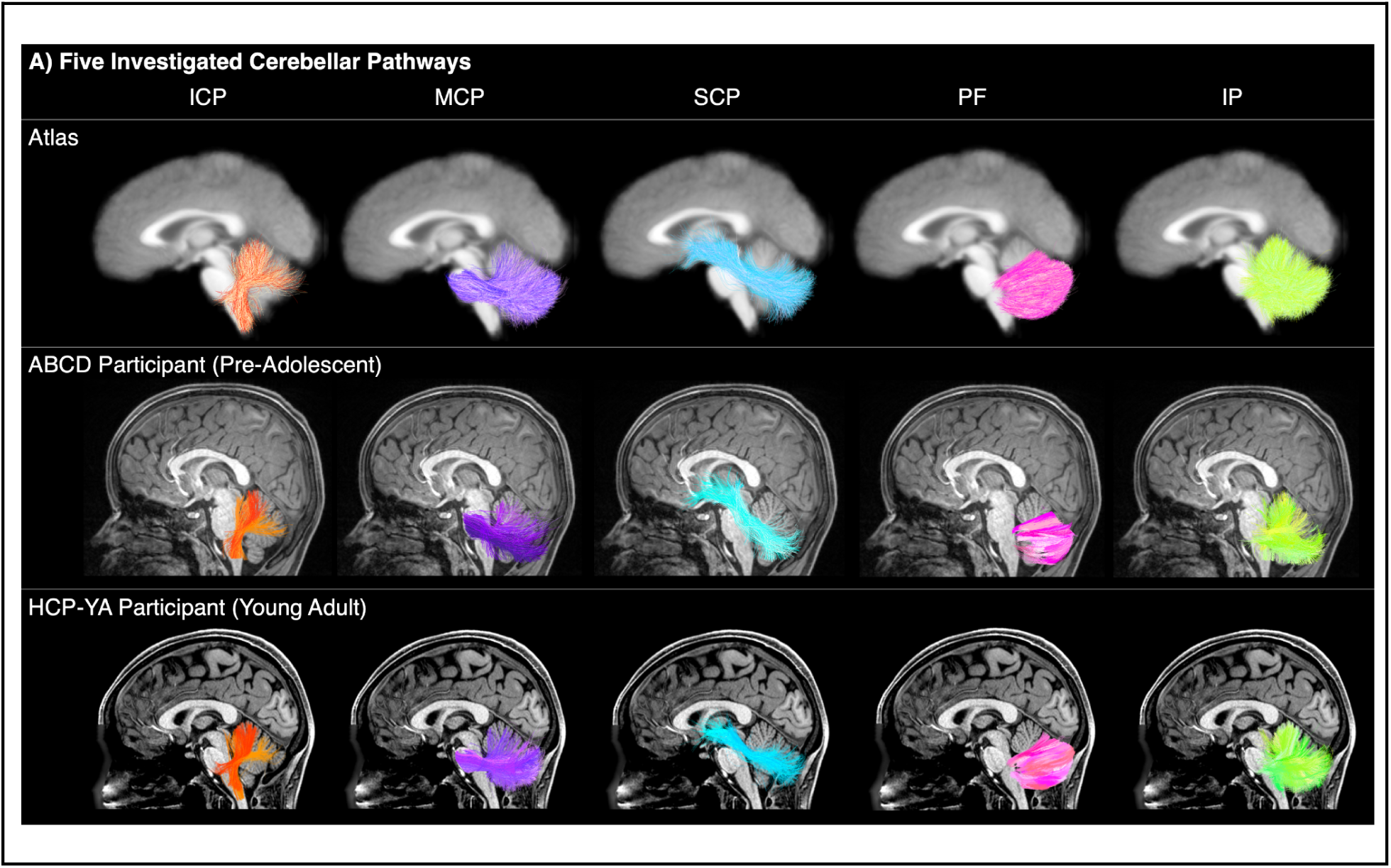
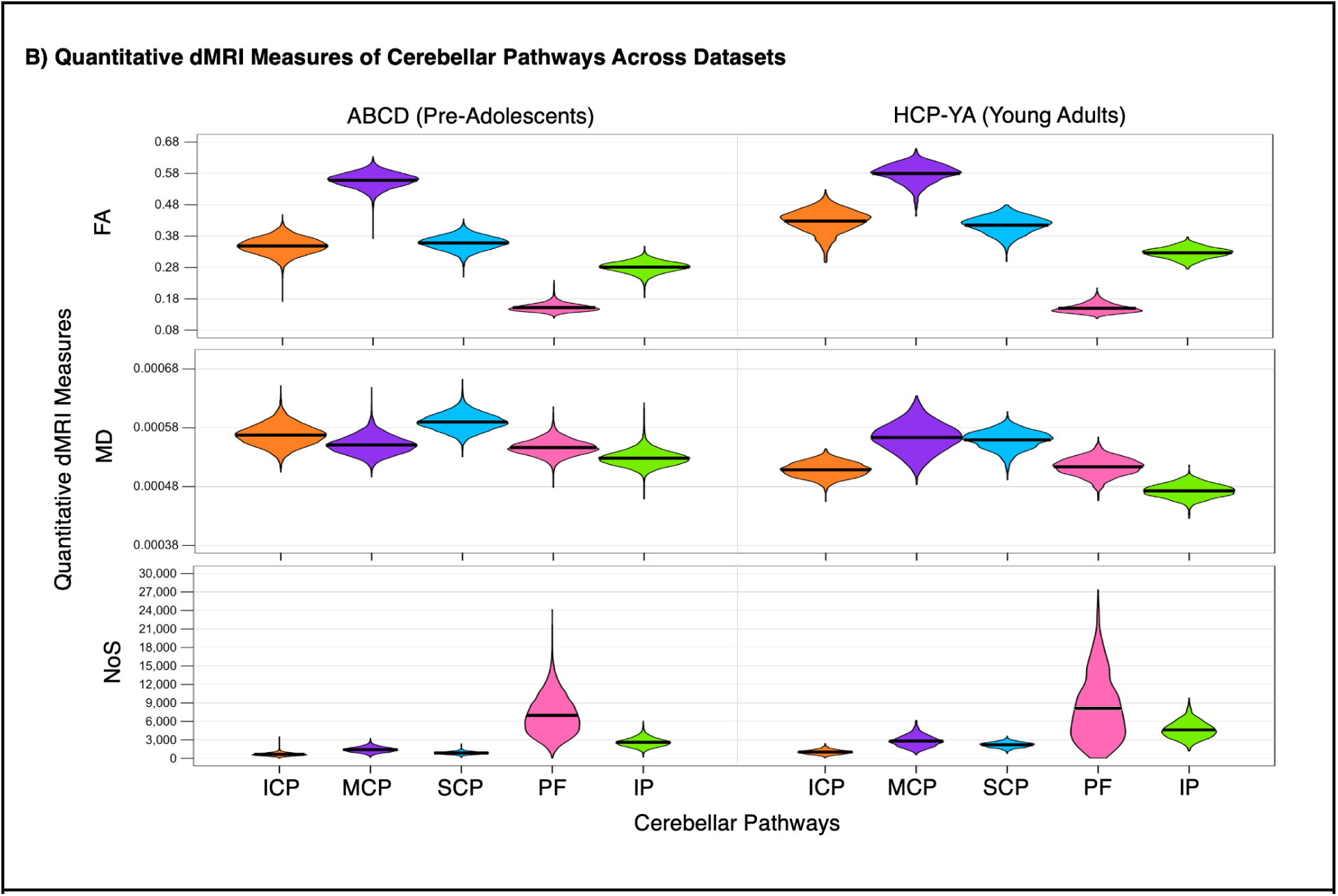
Consistent identification and quantification of the cerebellar pathways across datasets and age groups. **A)** Visualization of five cerebellar pathways in the atlas, a representative ABCD pre-adolescent participant, and a representative HCP-YA young adult. Pathways are shown from the left and overlaid on the corresponding atlas or participant T1-weighted image. **B)** Quantitative dMRI measures (FA, MD, and NoS) of cerebellar pathways show consistent distributions across datasets. Violin plots illustrate these measures for 8,848 pre-adolescents (ABCD) and 945 young adults (HCP-YA), with the black bar indicating the mean. Abbreviations: ICP, inferior cerebellar peduncle; MCP, middle cerebellar peduncle; SCP, superior cerebellar peduncle; PF, parallel fibers; IP, input and Purkinje fibers; ABCD, Adolescent Brain and Cognitive Development Study; HCP-YA, Human Connectome Project-Young Adult dataset; FA, fractional anisotropy; MD, mean diffusivity; NoS, Number of Streamlines.

Three dMRI measures—FA, MD, and NoS—are computed for each pathway (**Figure 1B**), representing diffusion anisotropy, diffusion magnitude, and structural connectivity strength^12,30^. Within each dataset, each pathway exhibits unique FA, MD, and NoS values, suggesting pathway-specific microanatomy characteristics (see **SI Table 1** for summary statistics). Similar distributions of dMRI measures across both datasets demonstrate consistent pathway identification and quantification. The mean FA of the cerebellar peduncles is higher in young adults (HCP-YA) than it is in pre-adolescent children (ABCD). Similarly, the mean MD of the cerebellar pathways is lower in adults than it is in pre-adolescent children, with the exception of MD of the MCP. In both datasets, we find the PF have the lowest FA and highest NoS, demonstrating our capacity to consistently quantify and effectively detect the PF.

### 2.2 Associations between cerebellar pathway measures & cognition in two datasets

We examine the relationship between cerebellar pathway measures and cognitive performance using multiple linear regression, with covariates including age, sex, and head motion (see Methods). We analyze seven assessments of cognitive performance from the NIH Toolbox Cognition Battery, including two measures of language, two measures of executive function, one measure of processing speed, and two measures of memory^31^ (see **SI Figure 1** and **SI Table 2** for distributions of measures).

In each dataset, 105 multiple linear regression models are created (5 pathways × 3 dMRI measures × 7 cognitive assessments) to evaluate the associations. Standardized *β* coefficients are used to estimate the strength and direction of each association on a uniform scale. Across both datasets, we find numerous significant associations between cerebellar pathway microstructure and cognitive performance (ABCD: 46 significant *β* coefficients; HCP-YA: 25 significant *β* coefficients). **Figure 2** provides a visualization of *β* coefficients across datasets, with violin plots showing significant *β* coefficient distributions after FDR correction. Results in **Figure 2A** (ABCD dataset) and **Figure 2B** (HCP-YA dataset) reveal variability in *β* coefficients across pathways, dMRI measures, and cognitive assessments. We observe larger significant relationships in the HCP-YA dataset (HCP-YA mean *β* = 0.12 vs. ABCD mean *β* = 0.06; *p* < 0.0001, t(28.38) = -4.78, 95% CI [-0.087, -0.035]) (**Figure 2D** and **SI Table 3**). In the HCP-YA dataset but not the ABCD dataset, we find one significant negative relationship between the PF and performance on a inhibitory control and attention assessment (Flanker) (**Figures 2B** and **2D**).

**Figure 2.**
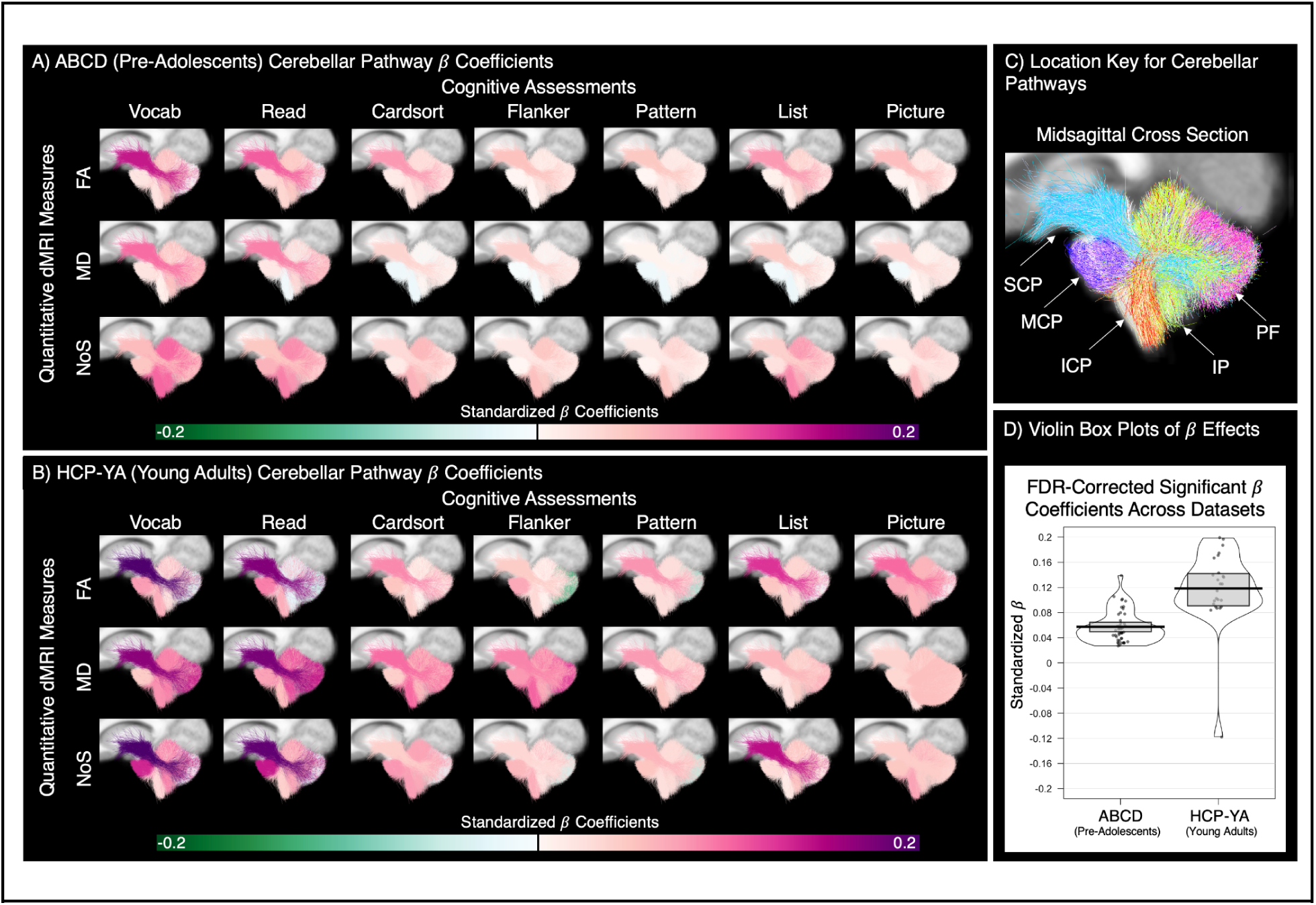
Associations between quantitative dMRI measures of the cerebellar pathways and performance on cognitive assessments. **(A, B)** Cerebellar pathways are colored based on *β* coefficients estimating relationships between dMRI measures and cognitive assessment performance in the ABCD **(A)** and HCP-YA **(B)** datasets. Rows represent three dMRI measures; columns show seven NIH Toolbox cognitive assessments. Pathways are displayed on a population mean T1w parasagittal slice (see pathway key in **C**). **(D)** Violin plots show distributions of significant *β* coefficients after FDR correction, with means (bold black lines) and 95% confidence intervals (gray boxes). Abbreviations: ICP, inferior cerebellar peduncle; MCP, middle cerebellar peduncle; SCP, superior cerebellar peduncle; PF, parallel fibers; IP, input and Purkinje fibers; ABCD, Adolescent Brain and Cognitive Development Study; HCP-YA, Human Connectome Project-Young Adult dataset; FA, fractional anisotropy; MD, mean diffusivity; NoS, Number of Streamlines; Vocab, NIH Toolbox Picture Vocabulary Test; Read, NIH Toolbox Oral Reading Recognition Test; Cardsort, NIH Toolbox Dimensional Change Card Sort Test; Flanker, NIH Toolbox Flanker Inhibitory Control and Attention Test; Pattern, NIH Toolbox Pattern Comparison Processing Speed Test; List, NIH Toolbox List Sorting Working Memory Test; Picture, NIH Toolbox Picture Sequence Memory Test.

The strongest associations involve the superior cerebellar peduncle (SCP) and the picture vocabulary and oral reading assessments (see **Figures 2A** and **2B**, and **Table 1** for more detail). In the ABCD dataset, the strongest association is between FA of the SCP and performance on the picture vocabulary assessment (*β* = 0.14, 95% CI [0.12, 0.16], p < 0.0001****). In the HCP-YA dataset, the strongest association is between NoS of the SCP and performance on the picture vocabulary assessment (*β* = 0.20, 95% CI [0.13, 0.26], p < 0.0001****). Notably, both picture vocabulary and oral reading assessments capture language-related abilities, suggesting a strong link between SCP microstructure and language performance across both adults and children.

**Table 1.** Consistently significant relationships between cerebellar pathway dMRI measures and cognitive measures in both the ABCD and HCP-YA datasets. Bolded and light blue rows indicate significant relationships between language measures (Read & Vocab) and SCP across all three dMRI measures in both datasets. The table reports FDR-corrected p-values and standardized β coefficients. Abbreviations: ICP, inferior cerebellar peduncle; MCP, middle cerebellar peduncle; SCP, superior cerebellar peduncle; PF, parallel fibers; IP, input and Purkinje fibers; ABCD, Adolescent Brain and Cognitive Development Study; HCP-YA, Human Connectome Project-Young Adult dataset; FA, fractional anisotropy; MD, mean diffusivity; NoS, Number of Streamlines; Vocab, NIH Toolbox Picture Vocabulary Test; Read, NIH Toolbox Oral Reading Recognition Test; Cardsort, NIH Toolbox Dimensional Change Card Sort Test; Flanker, NIH Toolbox Flanker Inhibitory Control and Attention Test; Pattern, NIH Toolbox Pattern Comparison Processing Speed Test; List, NIH Toolbox List Sorting Working Memory Test; Picture, NIH Toolbox Picture Sequence Memory Test.

|  |  | ABCD<br>(Pre-Adolescents) |  |  | HCP-YA<br>(Young Adults) |  |  |
| --- | --- | --- | --- | --- | --- | --- | --- |
| Cerebellar Pathway | Cognitive Assessment | $\beta$ | 95% CI | *p*-value | $\beta$ | 95% CI | *p*-value |
| **FA** |  |  |  |  |  |  |  |
| MCP | Read | 0.032 | [0.011, 0.053] | 0.0076** | 0.089 | [0.025, 0.15] | 0.032* |
| SCP | Cardsort | 0.077 | [0.056, 0.099] | < 0.0001**** | 0.084 | [0.02, 0.15] | 0.044* |
| SCP | List | 0.078 | [0.056, 0.099] | < 0.0001**** | 0.13 | [0.061, 0.19] | 0.0011** |
| SCP | Picture | 0.055 | [0.034, 0.077] | < 0.0001**** | 0.10 | [0.036, 0.16] | 0.013* |
| **SCP** | **Read** | **0.11** | **[0.084, 0.13]** | **< 0.0001\*\*\*\*** | **0.17** | **[0.11, 0.23]** | **< 0.0001\*\*\*\*** |
| **SCP** | **Vocab** | **0.14** | **[0.12, 0.16]** | **< 0.0001\*\*\*\*** | **0.20** | **[0.13, 0.26]** | **< 0.0001\*\*\*\*** |
| **MD** |  |  |  |  |  |  |  |
| SCP | Cardsort | 0.036 | [0.016, 0.057] | 0.0017** | 0.10 | [0.036, 0.16] | 0.013* |
| SCP | Flanker | 0.039 | [0.018, 0.06] | 0.00071*** | 0.087 | [0.024, 0.15] | 0.032* |
| **SCP** | **Read** | **0.089** | **[0.078, 0.12]** | **< 0.0001\*\*\*\*** | **0.17** | **[0.11, 0.24]** | **< 0.0001\*\*\*\*** |
| **SCP** | **Vocab** | **0.10** | **[0.068, 0.11]** | **< 0.0001\*\*\*\*** | **0.17** | **[0.1, 0.23]** | **< 0.0001\*\*\*\*** |
| PF | Read | 0.059 | [0.039, 0.08] | < 0.0001**** | 0.14 | [0.075, 0.2] | 0.00042*** |
| PF | Vocab | 0.058 | [0.038, 0.078] | < 0.0001**** | 0.14 | [0.072, 0.2] | 0.00037*** |
| IP | Read | 0.032 | [0.012, 0.053] | 0.0056** | 0.11 | [0.049, 0.18] | 0.0042** |
| IP | Vocab | 0.055 | [0.033, 0.074] | < 0.0001**** | 0.089 | **[0.024, 0.15]** | 0.032* |
| **NoS** |  |  |  |  |  |  |  |
| MCP | Read | 0.062 | [0.042, 0.083] | < 0.0001**** | 0.14 | [0.075, 0.2] | 0.00027*** |
| MCP | Vocab | 0.048 | [0.028, 0.068] | < 0.0001**** | 0.13 | [0.069, 0.19] | 0.00037*** |
| SCP | List | 0.039 | [0.018, 0.06] | 0.00082*** | 0.14 | [0.077, 0.21] | 0.00027*** |
| **SCP** | **Read** | **0.047** | **[0.027, 0.068]** | **< 0.0001\*\*\*\*** | **0.19** | **[0.12, 0.25]** | **< 0.0001\*\*\*\*** |
| **SCP** | **Vocab** | **0.056** | **[0.035, 0.076]** | **< 0.0001\*\*\*\*** | **0.20** | **[0.13, 0.26]** | **< 0.0001\*\*\*\*** |
| IP | Vocab | 0.10 | [0.079, 0.12] | < 0.0001**** | 0.087 | [0.02, 0.15] | 0.044* |

### 2.3 Across-dataset comparison of the relationship between cerebellar pathway measures & cognition

We compare associations between cerebellar pathway measures and cognition in pre-adolescent children (ABCD) and young adults (HCP-YA). Of the 25 significant relationships in the HCP-YA dataset, 20 are also significant in the ABCD dataset, primarily involving SCP measures and language assessments (**Table 1**). These 20 shared relationships span FA, MD, and NoS measures of the MCP, SCP, IP, and PF, and are linked to language, memory, and executive function. Notably, 14 of these 20 relationships are with language assessments, and 12 specifically involve the SCP. Significant relationships between the SCP and language measures (Picture Vocabulary & Oral Reading) are observed across all three dMRI measures in both datasets (see bolded and highlighted rows in **Table 1**). This pattern suggests that many significant relationships between dMRI measures and cognition observed in young adults are also present in children, particularly those involving SCP measures and language performance.

Next, we assess the consistency of cerebellar dMRI-cognition relationships across pre-adolescents and young adults by evaluating the correlation of *β* coefficients between the two datasets. A moderate and significant correlation is found between the 105 *β* coefficients from the ABCD and HCP-YA datasets (Pearson’s *r* = 0.47, 95% CI [0.30, 0.60], *p* < 0.0001) (**SI Figure 2**), demonstrating consistency of overall results across age groups.

We then analyze the consistency of results for each dMRI measure. **Figure 3** presents the correlation of standardized *β* coefficients across datasets for FA, MD, and NoS. FA and MD *β* coefficients show strong and significant correlations across datasets (FA: *r* = 0.67, 95% CI [0.44, 0.82], *p <* 0.0001; MD: *r* = 0.75, 95% CI [0.56, 0.87], *p* < 0.0001). This finding demonstrates consistent relationships between cerebellar pathway microstructure (FA and MD) and cognition in both pre-adolescent children and young adults.

**Figure 3.**
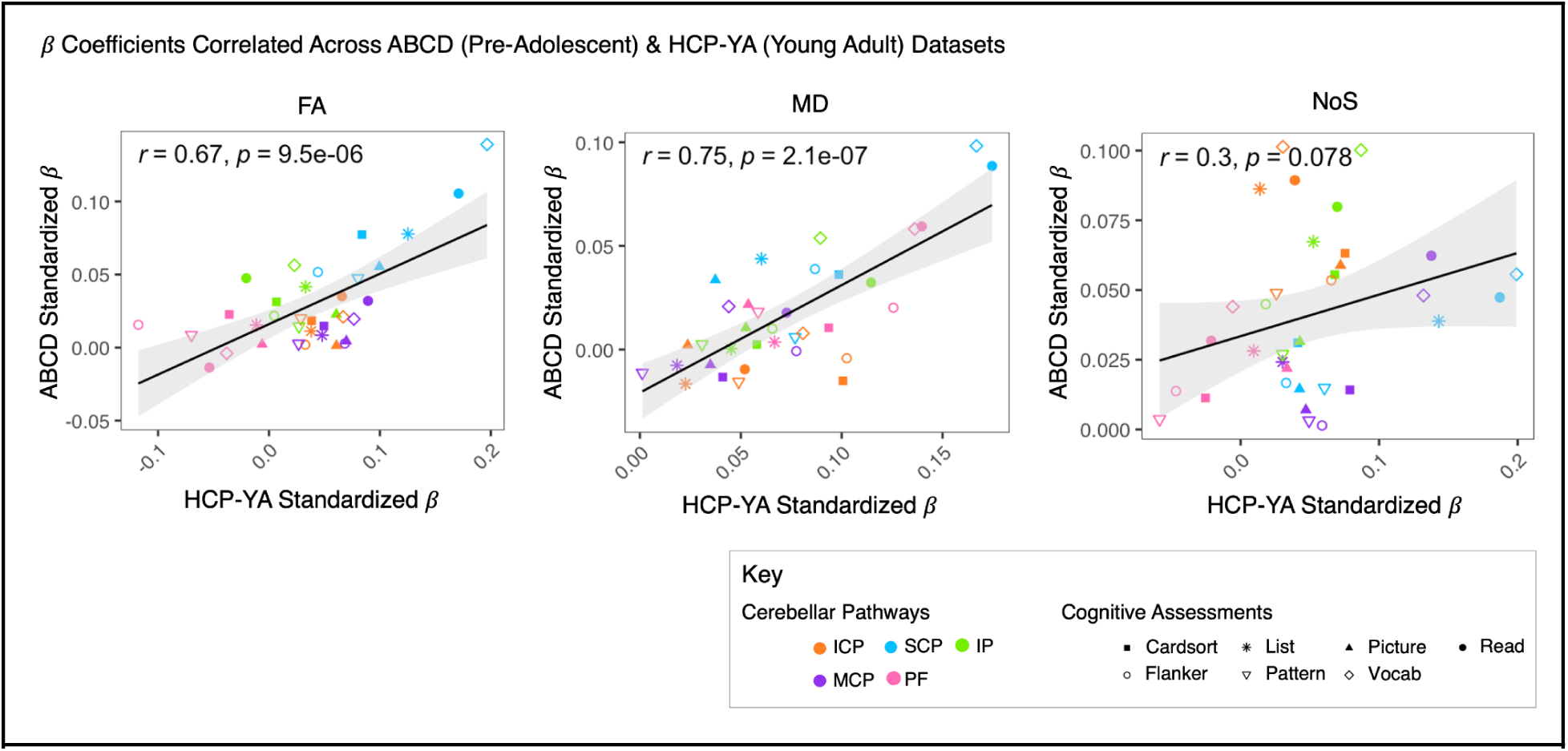
Correlation of standardized *β* coefficients across ABCD (pre-adolescent) and HCP-YA (young adult) datasets. Scatterplots illustrate correlations of standardized *β* coefficients between the ABCD (pre-adolescent) and HCP-YA (young adult) datasets for FA (left), MD (middle), and NoS (right). Each scatterplot includes 35 points (7 cognitive assessments × 5 cerebellar pathways). *β* coefficients from the HCP-YA dataset are plotted on the x-axis, and those from the ABCD dataset are on the y-axis. Each point represents data from over 9,000 participants, with colors denoting cerebellar pathways and shapes representing cognitive assessments (see **Key**). Gray shaded region shows 95% confidence interval. Abbreviations: ICP, inferior cerebellar peduncle; MCP, middle cerebellar peduncle; SCP, superior cerebellar peduncle; PF, parallel fibers; IP, input and Purkinje fibers; ABCD, Adolescent Brain and Cognitive Development Study; HCP-YA, Human Connectome Project-Young Adult dataset; FA, fractional anisotropy; MD, mean diffusivity; NoS, Number of Streamlines; Vocab, NIH Toolbox Picture Vocabulary Test; Read, NIH Toolbox Oral Reading Recognition Test; Cardsort, NIH Toolbox Dimensional Change Card Sort Test; Flanker, NIH Toolbox Flanker Inhibitory Control and Attention Test; Pattern, NIH Toolbox Pattern Comparison Processing Speed Test; List, NIH Toolbox List Sorting Working Memory Test; Picture, NIH Toolbox Picture Sequence Memory Test.

In contrast, NoS *β* coefficients have a weaker and non-significant correlation (*r* = 0.3, *p* = 0.078). Across datasets, we observe notable differences in the relationships between NoS and cognitive performance, particularly in the cerebellar peduncles. First, significant relationships between cognition and NoS of the SCP and MCP exhibit substantially larger *β* coefficients (more than double) in the HCP-YA dataset compared to the ABCD dataset (**Table 1**). Second, in ABCD, NoS in the ICP is significantly associated with all seven cognitive assessments—especially Vocab, Read, and List—but shows no significant relationship in HCP-YA (**Figure 2A** **& 2B, SI Table 3**).

## 3. Discussion

This study investigates the relationships between dMRI measures of cerebellar pathways and cognitive performance, revealing numerous significant associations that are consistent across two large datasets of pre-adolescent children (ABCD) and young adults (HCP-YA). By leveraging data from over 9,000 participants and employing consistent tractography methods and assessment measures, we provide strong evidence of the cerebellum’s contribution to cognition. These findings, alongside prior research^32^, underscore the importance of large neuroimaging datasets in advancing our understanding of brain-cognition relationships.

Our findings demonstrate striking consistency despite notable differences in dataset sample size, resolution, and participant ages. First, the ABCD dataset has over 8,000 participants compared to 945 participants in HCP-YA: the stronger statistical power may explain the greater number of significant *β* coefficients found in the ABCD dataset (46 vs. 25). Second, the higher dMRI resolution of the HCP-YA data (1.25 mm isotropic vs. 1.7 mm isotropic in ABCD)^33,34^ improves pathway delineation and reduces partial volume effects. This may influence the higher FA in the cerebellar peduncles (**Figure 1B**) and the larger associations in HCP-YA (**Figure 2D**) as observed in previous research^32^. Finally, maturational differences between pre-adolescent children (ABCD) and young adults (HCP-YA) may further explain variations in significance and effect size, as cerebellar and cognitive development continue through adolescence^35–39^.

While each of these differences have the potential to influence our findings, our results demonstrate multiple layers of consistency. First, cerebellar pathways were reliably identified and quantified using the same neuroanatomical fiber atlas across datasets (**Figures 1A** and **1B**). Second, in both datasets, dMRI measures of the cerebellar pathways are significantly associated with performance on cognitive assessments, with many findings showing significance across datasets (Figure 2 and **Table 1**). Third, our findings are strongly and significantly correlated across datasets, as shown in **Figure 3** and visually evident in **Figures 2A** and **2B**. Overall, these findings point to consistent associations between dMRI measures of the cerebellar pathways and performance on cognitive assessments in pre-adolescents and young adults.

Among the significant findings, we identified strong associations between SCP dMRI measures (FA, MD, NoS) and language performance (Vocab & Read) that were consistent across both datasets (**Figure 2**, **Table 1**). These dMRI measures are not orthogonal (**SI Figure 3**), suggesting they may reflect common underlying microstructural components. Similarly, strong correlations between vocabulary and reading performance in both the ABCD and HCP-YA datasets (Vocab & Read, **SI Figure 4**) suggest a shared cognitive construct, such as language or “crystalized intelligence”^40,41^. Together, these findings suggest a stable link between SCP microstructure and accumulated language knowledge, consistent with prior research that links the SCP to word reading and verbal fluency^14,25^. As the cerebellum’s primary output pathway to the cerebrum, the SCP likely plays a key role in cerebellar contributions to language processing. Future studies may further investigate the SCP’s role in language disorders and its broader contributions to development and language-related cognition.

Unlike the microstructure measures (FA, MD), the *β* coefficients related to the connectivity measure (NoS) are not significantly correlated across datasets (Figure 3). While this finding may be influenced by acquisition differences or other differences across datasets discussed above, it may also indicate that the NoS measure is more sensitive to developmental differences related to cognition. This interpretation aligns with other evidence suggesting that, compared to FA and MD, NoS is a more predictive measure of both cognitive performance and age^42,43^.

One remarkable feature of the NoS measure for the language assessments (Vocab & Read, **Figure 2**) is the developmental shift in cerebellar pathway emphasis. In children, the ICP is strongly weighted, but by young adulthood, this emphasis diminishes as the SCP and MCP gain prominence. This observation may align with findings that earlier cerebellar damage or surgery is associated with poorer long-term functional recovery^44^, and is a potentially impactful area for future research. It can be hypothesized that the early emphasis on the ICP reflects the development of climbing fiber inputs on Purkinje cells.

Climbing fibers from the inferior olivary nucleus, which are conveyed through the ICP and are responsible for the powerful and electrophysiologically identifiable complex spikes in Purkinje cells, have considerable theoretical importance for providing teaching signals that modify parallel fiber/Purkinje cell synapses^45,46^, thereby providing a hypothetical learning mechanism that has received support from experimental studies^47^. By young adulthood, the shift towards the SCP and MCP may indicate the maturation of cerebro-cerebellar closed-loop pathways described by Strick and colleagues^48^ involve cerebellar efferents projecting to neocortical regions via dentato-thalamo-cortical projections through the SCP, and afferents from those same cortical regions via cortico-ponto-cerebellar projections through MCP. Overall, the significant associations of cerebellar pathways with language performance aligns with functional neuroimaging studies demonstrating cerebellar contributions to reading^49^ and its responsiveness to both phonological^50,51^ and semantic aspects^51–53^ of language processing aspects integral to the Vocab & Read assessments.

A similar trend in ICP emphasis for the NoS measure is seen in the List assessment, which is an executive working memory task that also exhibited strong association between cerebellar pathways and task performance. Cerebellar activation has been found consistently in verbal working memory functional neuroimaging studies^2,54,55^. Like the language assessments, ICP strength in the List assessment is prominent in pre-adolescents and diminishes in young adulthood, replaced by strong weighting of SCP. In contrast to the language assessments, the NoS measure for MCP on the List task is not strongly weighted in young adulthood for working memory as much as it is in language assessments. This perhaps reflects the fact that the language assessments involve processing of well-learned stimuli whereas the working memory assessment involves processing of novel sequences of words that might benefit more from cerebellar forward model predictions^56^ through the SCP than from feedback via MCP.

Additionally, our dMRI tractography analysis identified horizontal streamlines in the cerebellar cortex resembling parallel fibers (**Figure 1A, SI Figure 5**). The granule cells, the most abundant neurons in the mammalian brain^57^, form a dense layer of predominantly unmyelinated axons, known as the parallel fibers, which extend laterally across the molecular layer of the cerebellar cortex^58^. Consistent with these characteristics, we observed that PF exhibits the highest NoS and lowest mean FA (**Figure 1B**), potentially related to their numerosity and unmyelinated structure. Previous studies have detected PF using dMRI^9,10^, and our investigation may provide new insights into their microstructural anatomy and cognitive relevance.

We find that MD of PF is significantly associated with vocabulary and reading performance in both the ABCD and HCP-YA datasets (**Figures 2A** and **2B**, **Table 1**), suggesting a consistent relationship between PF microstructure and language function across age groups. Additionally, FA of the PF is significantly associated with inhibitory control (NIH Toolbox Flanker Test) in young adults (HCP-YA) but not in pre-adolescents (ABCD) (**Figure 2B**, **SI Table 3**), suggesting a role of the PF in inhibitory control that may develop with age. This finding is particularly intriguing and may reflect broader cerebellar contributions to the regulation of attentional and error-monitoring mechanisms^59,60^. Previous work investigating the cognitive role of the cerebellar granule cells in mice has associated the granule cells with reward anticipation^61^. However, how these findings in rodents may translate to human cognitive functions like language and inhibitory control remains unclear. Future research may continue to investigate the links between reward anticipation, inhibitory control, attention, and language in both animal models and humans.

Overall, this exploratory study establishes a foundation for investigating the relationships between cerebellar pathway microstructure and cognition. The consistency of our results across datasets with varying sample sizes, resolutions, and participant ages suggests that these relationships are likely generalizable and replicable in other datasets. Future studies may continue to build on these findings by exploring similar associations in other large, open-access dMRI datasets that include cognitive performance measures, and by leveraging complex models to explore non-linear relationships and mediation effects. Future work may also leverage new, high-resolution dMRI acquisitions to create more detailed atlases of cerebellar connections, enabling assessment of potential lateralized and localized effects. While we identified significant associations, our findings do not establish causal relationships between measures of the cerebellar pathways and cognitive performance. Future studies may explore interventional effects or causal inference methodology to potentially shed light on causal aspects of these relationships.

Here, we focused on large, non-clinical populations of pre-adolescents and young adults, which enabled us to identify significant relationships between dMRI measures of cerebellar pathways and cognitive performance. However, these findings may not fully capture the anatomical and cognitive variability across the human lifespan and in clinical populations. Previous research has investigated dMRI measures of the cerebellar pathways across neurodevelopmental, neurodegenerative, and cognitive disorders - including attention-deficit hyperactivity disorder, Parkinsonian syndromes, schizophrenia, cerebellar mutism, and ataxia^20,22,23,62^. Future research may continue to investigate the cerebellar pathways in relation to cognitive function across such conditions to further inform diagnosis, treatment, and intervention strategies. By establishing baseline relationships between cerebellar pathways and cognition in healthy young adults and pre-adolescent children, this study provides a foundation for targeted hypotheses about the cerebellum’s role in various disorders and developmental time points.

In conclusion, using two separately acquired large population datasets of young adults (HCP-YA) and preadolescents (ABCD), we find significant relationships between quantitative dMRI measures of the cerebellar pathways and performance on cognitive assessments. In both datasets, we find that the strongest and most consistently significant relationships exist between the dMRI measures of the superior cerebellar peduncle (SCP) and performance on language assessments, suggesting that the SCP plays an important role in language function. In the adult dataset, we demonstrate a significant relationship between quantitative dMRI measures of the parallel fibers (PF) and performance on an inhibitory control assessment, suggesting that the parallel fibers, the most numerous neurons in the human brain, may play an important role in attentional control. Overall, we find numerous significant associations and results that correlate across age groups, strongly and consistently implicating the cerebellar pathway microstructure in cognition.

## 4. Materials and Methods

### 4.1 HCP-YA & ABCD Datasets

This study investigates relationships between cerebellar pathways and cognitive performance using data from two large openly released neuroimaging datasets: the HCP-YA dataset^63^ and the ABCD dataset^29^. The HCP-YA dataset includes over 1,000 healthy young adults, while the ABCD dataset includes over 10,000 pre-adolescent children. We utilize the baseline data from the ABCD dataset, which is longitudinal with multiple data collection timepoints. Access to these datasets was granted by the Human Connectome Project and the National Institute of Mental Health Data Archive (NDA). The study protocol was approved by the Mass General Brigham Institutional Review Board.

### 4.2 Inclusion & Exclusion Criteria

This study utilizes the largest possible set of participants from the ABCD and HCP-YA with high-quality dMRI scans and complete cognitive performance data. We use study-specific quality control (QC) data to determine inclusion and exclusion criteria for these datasets (see **SI Figure 6** for the inclusion process).

In the ABCD dataset, some dMRI scans lacked full inferior brain coverage, occasionally omitting parts of the cerebellum^64^. Field-of-view cutoffs were assessed using both automated and manual methods^65^. To balance sample size and data quality, we excluded participants with more than 30% intersection of the brain mask with the ventral frame border (apqc_dmri_fov_cutoff_ventral < 30, n=126), those with a manual cutoff rating above zero (dmri_dti_postqc_qucutoff > 0, n=87), those flagged for poor post-processing quality (dmri_dti_postqc_qc=1, n=122), those with undetected cerebellar pathways (n=1), and those missing any of the seven NIH Toolbox Cognitive Battery measures (n=161). The final ABCD sample included 8,848 participants aged 8 years 11 months to 11 years 1 month (mean: 9 years 11 months; 4,606 male, 4,242 female).

The HCP-YA dataset comprises healthy young adults aged 22 to 35 from Missouri, USA^63^, where ‘healthy’ was broadly defined, encompassing individuals with a history of smoking, overweight status, or recreational drug and alcohol use. The dataset also includes siblings, including twins^63^. Of the initial 1,206 participants, 1,065 had preprocessed dMRI data. We excluded those with QC issues due to anatomical anomalies or head coil instability (QC_Issue = A or C; n=44 and n=73, respectively)^66^, as well as participants missing any of the seven NIH Toolbox measures (n=3). The final HCP-YA sample included 945 participants aged 22 to 37 years (mean: 28 years 9 months; 518 male, 427 female). All five cerebellar pathways of interest were detected in these participants.

### 4.3 NIH Toolbox Cognition Battery Assessments

Both HCP-YA and ABCD datasets include seven individual-level cognitive assessments from the NIH Toolbox Cognition Battery. These assessments measure executive function, attention, episodic memory, language, processing speed, and working memory^39,67^ and comprise the Dimensional Change Card Sort, Flanker Inhibitory Control and Attention, List Sorting Working Memory, Oral Reading Recognition, Pattern Comparison Processing Speed, Picture Sequence Memory, and Picture Vocabulary tests^68^. The NIH Toolbox Cognition Battery is validated in both children and adults^68,69^. We use uncorrected scores—reflecting performance relative to the entire NIH Toolbox normative sample—without adjusting for age, sex, or other factors^70,71^. Uncorrected scores enable us to account for effects of age and sex demographics unique to our participant samples by adding them as covariates in our regression analyses. Additionally, uncorrected scores can capture the maturational differences in cognitive performance between pre-adolescents (ABCD) and young adults (HCP-YA) (**SI Figure 1**). Higher uncorrected scores indicate better performance. See **SI Figure 4** and **SI Table 2** for measures and summary statistics.

### 4.4 dMRI Acquisition, Pre-processing, and ABCD dMRI Harmonization

The investigated ABCD participants were scanned at 18 sites across the USA using 3T Siemens and GE scanners with varying software versions^34^. ABCD dMRI scans of these participants used multiband EPI (slice acceleration factor=3, 1.7 mm isotropic resolution) with 96 directions, seven b=0 volumes, four b-values (b=500, 1000, 2000, 3000 s/mm²) with 6, 15, 15, and 60 directions respectively, and TR/TE 4100/88 ms (Siemens) and 4100/81.9 ms (GE)^34^. The HCP-YA data, in contrast, were collected at a single site using a single specialized 3T Siemens scanner^72^. HCP-YA dMRI scans used multiband EPI (slice acceleration factor=3, 1.25 mm isotropic resolution) with 270 directions, eighteen b=0 volumes, and three b-values (b= 1000, 2000, 3000 s/mm²) with 90 directions each, collected with TR/TE=5520/89.5 ms.

Both datasets underwent minimal pre-processing (eddy current, motion, and distortion correction and isotropic resampling)^34,73^. To reduce scanner-, protocol-, and site-related biases in ABCD data, we used harmonized ABCD dMRI data^27^ using a rotation-invariant spherical harmonics approach that removes scanner-related signal biases while preserving inter-subject variability^27,74^. ABCD participants scanned on Philips scanners were excluded from the harmonized ABCD dMRI dataset^27^ used in the present study (**SI Figure 6**) due to ringing artifacts, motion, excessive smoothing, and inconsistencies in baseline acquisitions. The harmonized baseline ABCD data are accessible through the NDA (https://nda.nih.gov/), with instructions on how to download them provided in our previous work^27,74^.

### 4.5 Whole Brain UKF Tractography

Whole-brain tractography was previously conducted on the study sample, which includes 8,848 participants from the ABCD dataset and 945 participants from the HCP-YA dataset^27,75^. From the minimally pre-processed HCP and harmonized ABCD dMRI data, we selected the b=3000 shell and b=0 volumes, as the b=3000 shell is optimal for resolving crossing fibers and represents the most comparable acquisition between ABCD and HCP^27,75–77^. Tractography was performed using the Unscented Kalman Filter (UKF) tractography method^78,79^ a multi-tensor approach implemented in ukftractography (github.com/pnlbwh/ukftractography) that can estimate fiber-specific microstructure measures. UKF tractography reliably identifies cerebellar pathways across the lifespan^11^. Quality control of the tractography was performed visually and quantitatively using whitematteranalysis (WMA) (github.com/SlicerDMRI/whitematteranalysis).

### 4.6 Identification of Cerebellar Pathways & Measure Extraction

Whole-brain tractography in the ABCD and HCP-YA datasets was previously parcellated using WMA^27,75^, which applies a neuroanatomically curated pathway atlas^11^. This approach consistently identifies pathways across the lifespan, health conditions, and acquisitions with high test-retest reliability^11,80^. The atlas parcellates whole-brain tractography into fiber clusters, which have been curated by a neuroanatomist into recognized cerebellar pathways (**SI Table 4**). Five cerebellar pathways are examined: the inferior (ICP), middle (MCP), and superior (SCP) cerebellar peduncles, input & Purkinje fibers (IP), and parallel fibers (PF) (**Figure 1**, **SI Figure 5**). All pathways are examined bilaterally to avoid bisecting decussating fibers. For each individual’s cerebellar pathways, we compute dMRI measures of fractional anisotropy (FA), mean diffusivity (MD), and number of streamlines (NoS), which are widely used to study brain microstructure, cognition, and development^12^. FA and MD are averaged across all streamline points in a pathway, while NoS is the total number of streamlines in a pathway^12^.

### 4.7 Statistical Analyses & Visualizations

Multiple linear regression models were used to examine the association between pathway microstructure and cognitive performance. Linear models were chosen to ease scientific interpretation and provide baseline relationships to support future research. Following previous work, in the multiple regression models, a cerebellar pathway dMRI measure (FA, MD, NoS) was considered as dependent variable and a cognitive performance score was considered as independent variable^75^. Regression models were fitted separately for ABCD and HCP-YA. In HCP-YA models, we adjusted for age and sex^75^; in ABCD, we adjusted for age, sex, and head motion (dmri_meanmotion), given higher motion in children^81^. In total, 210 models were run (three microstructural measures × five pathways × seven tests × two datasets). We extract p-values and *β* coefficients, apply Benjamini & Hochberg false discovery rate (FDR) correction (across 105 models for each dataset)^82^, and consider FDR-adjusted p<0.05 significant. Standardized *β* coefficients are computed using “lm.beta”^83^. We use Welch’s t-tests to compare mean standardized *β* across datasets and Pearson’s correlation to compare *β* values from ABCD and HCP-YA. All analyses were exploratory and conducted in R^84^. Plots were generated using the packages “ggplot2” and “yarrr”^85,86^. Cerebellar visualizations are created in 3D Slicer with SlicerDMRI^87^.

## Data and Code Availability

The data used in this project are from the Adolescent and Cognitive Development Study (ABCD) dataset, accessible via the National Institute of Mental Health Data Archive (NDA) at nda.nih.gov/abcd, and the Human Connectome Project Young Adult (HCP-YA) dataset, available through the ConnectomeDB at db.humanconnectome.org. Additionally, we leveraged the harmonized ABCD data, available via the NDA (see ^27,88^). The code used for data analysis will be available upon publication at github.com/lzekelma/cerebellar-pathway-cognition-9000.

## Acknowledgments & Funding Sources

This work was supported by the National Institutes of Health (NIH) grants: R01MH132610, R01MH125860, R01MH119222, R01MH116173, R01NS125307, R01NS125781, R01MH112748, R01AG042512, R01MH128278, P41EB015902, T32DC000038, and K24MH116366; the National Key R&D Program (No. 2023YFE0118600); the National Natural Science Foundation (No. 62371107); Brigham and Women’s Hospital Department of Radiology Sundry Fund; Brigham and Women’s Hospital Program for Interdisciplinary Neurosciences (Lawrence & Tina Rand); Brain & Behavior Research Foundation NARSAD Young Investigator Award.

Data used in the preparation of this article were obtained from the Adolescent Brain Cognitive Development (ABCD) Study (https://abcdstudy.org), held in the NIMH Data Archive (NDA). This is a multisite, longitudinal study designed to recruit more than 10,000 children aged 9-10 and follow them over 10 years into early adulthood. The ABCD Study® is supported by the National Institutes of Health and additional federal partners under award numbers U01DA041048, U01DA050989, U01DA051016, U01DA041022, U01DA051018, U01DA051037, U01DA050987, U01DA041174, U01DA041106, U01DA041117, U01DA041028, U01DA041134, U01DA050988, U01DA051039, U01DA041156, U01DA041025, U01DA041120, U01DA051038, U01DA041148, U01DA041093, U01DA041089, U24DA041123, U24DA041147. A full list of supporters is available at https://abcdstudy.org/federal-partners.html. A listing of participating sites and a complete listing of the study investigators can be found at https://abcdstudy.org/consortium_members/. ABCD consortium investigators designed and implemented the study and/or provided data but did not necessarily participate in the analysis or writing of this report. This manuscript reflects the views of the authors and may not reflect the opinions or views of the NIH or ABCD consortium investigators. The ABCD data repository grows and changes over time. The ABCD data used in this report came from DOI: 10.15154/1520591 (Data Release 3.0).

Human Connectome Project Young Adult (HCP-YA) data were provided, in part, by the Human Connectome Project, WU-Minn Consortium (Principal Investigators: David Van Essen and Kamil Ugurbil; 1U54MH091657) funded by the 16 NIH Institutes and Centers that support the NIH Blueprint for Neuroscience Research; and by the McDonnell Center for Systems Neuroscience at Washington University.

## Author Contributions

LRZ - designed research, performed research, analyzed data, wrote paper; SCK - performed research, analyzed data, wrote paper; YC - performed research, analyzed data, wrote paper; MA - performed research, analyzed data, wrote paper; JHL - performed research, wrote paper; JR - performed research, wrote paper; SP - performed research, wrote paper; ZL - performed research, analyzed data, wrote paper; JED - performed research, wrote paper; LCB - performed research, wrote paper; NM - performed research, wrote paper; YR - performed research, wrote paper; FZ - performed research, analyzed data, wrote paper; AJG - designed research, performed research, wrote paper; LJO - designed research, performed research, wrote paper.

