## Supplementary Information for "Consistent cerebellar pathway-cognition associations across pre-adolescents & young adults: a diffusion MRI study of 9000+ participants"

### SI Figures

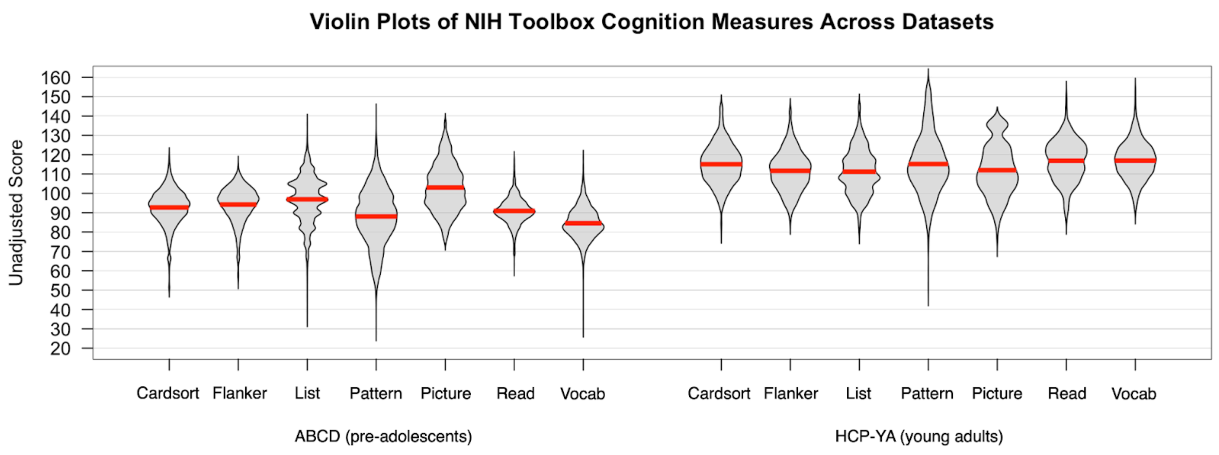


**SI Figure 1.** Distribution of unadjusted performance scores on NIH Toolbox Cognition Battery across datasets. Individual participant cognitive performance scores were included in both datasets. Unadjusted scores were normed without taking into account age and gender[1]. In these datasets, the NIH Toolbox Cognition Battery includes seven assessments that measure different cognitive abilities: the Dimensional Change Card Sort Test (Cardsort; executive function, shifting), Flanker Inhibitory Control and Attention Test (Flanker; executive function, attention), List Sorting Working Memory Test (List; memory, working), Oral Reading Recognition Test (Read; language, expressive), Pattern Comparison Processing Speed Test (Pattern; processing speed), Picture Sequence Memory Test (Picture; memory, episodic), and the Picture Vocabulary Test (Vocab; language, receptive)[2].

Note on NIH Toolbox Cognition Battery platform administration: NIH Toolbox Cognition Battery was administered via a web platform to participants in the HCP-YA dataset and via an app platform to the participants in the ABCD dataset. Uncorrected scores from all assessments are understood to be comparable across platforms with the exception of scores from the Picture Sequence Memory Test [3]. Thus, cross-dataset comparisons relating to performance on the Picture Sequence Memory Tests should be interpreted with additional caution. Picture Sequence Memory Test is included our analysis in order to have a measure of episodic memory function and in the interest of analyzing a more complete battery of assessments.

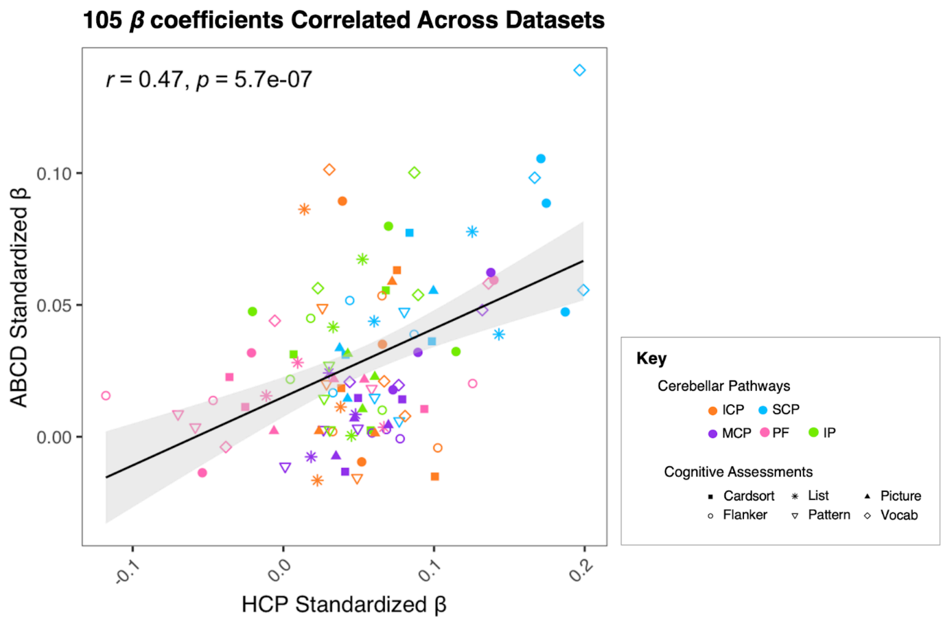


**SI Figure 2.** Correlation of standardized $\beta$ coefficients across ABCD (pre-adolescent) and HCP-YA (young adult) datasets. Scatterplots illustrate correlations of standardized $\beta$ coefficients between the ABCD (pre-adolescent) and HCP-YA (young adult) datasets. Each scatterplot includes 105 points (7 cognitive assessments × 5 cerebellar pathways × 3 dMRI measures). $\beta$ coefficients from the HCP-YA dataset are plotted on the x-axis, and those from the ABCD dataset are on the y-axis. Each point represents data from over 9,000 participants, with colors denoting cerebellar pathways and shapes representing cognitive assessments (see **Key**). Gray shaded region shows 95% confidence interval.

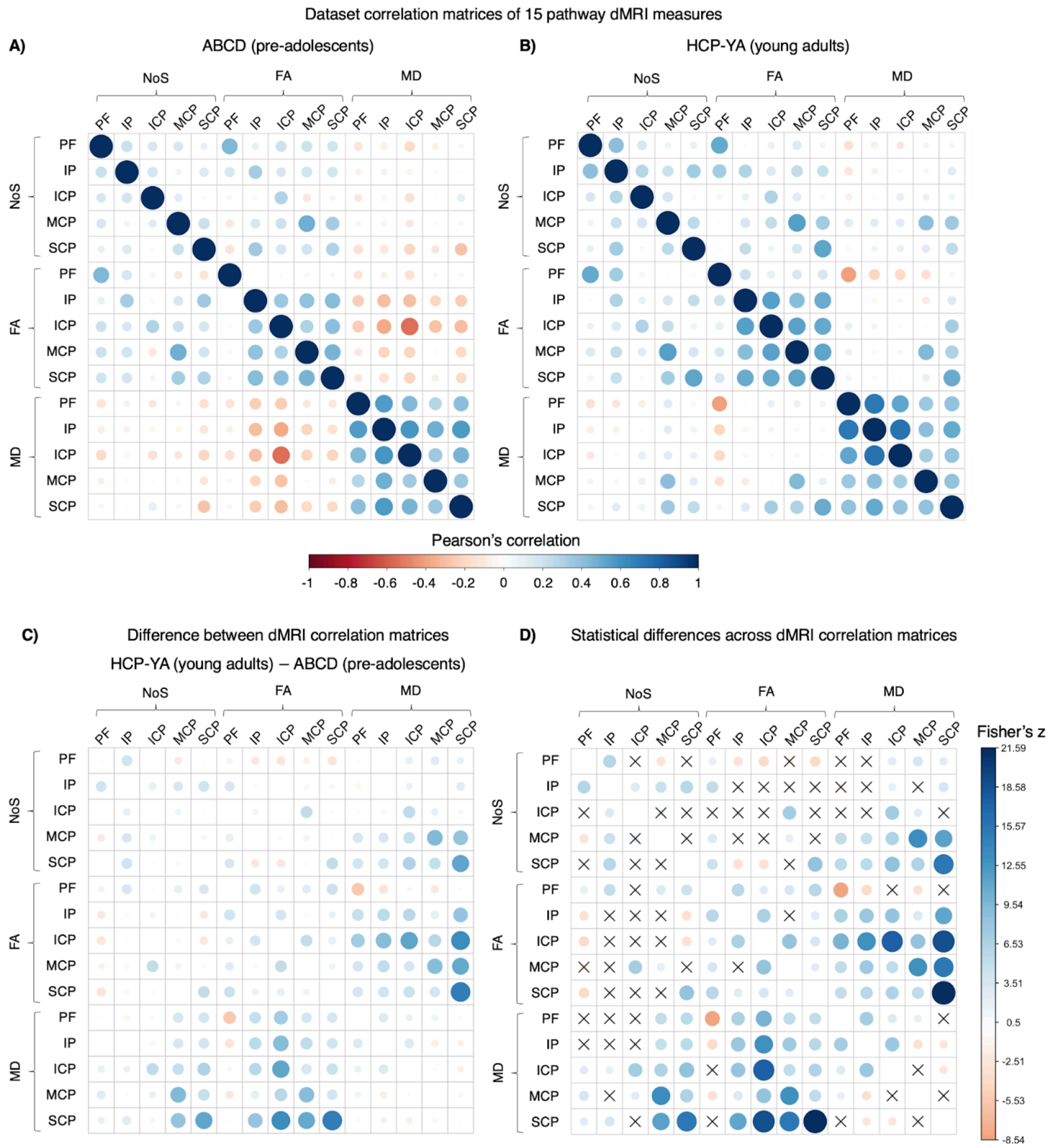


**SI Figure 3.** Dataset correlation matrices of dMRI measures. Correlation matrices of 15 pathway dMRI measures (5 pathways x 3 dMRI measures) in **(A)** the ABCD dataset and **(B)** the HCP-YA dataset. Correlations were calculated using Pearson's correlation coefficient and plotted using the corrplot package in R[4]. Color indicates direction and magnitude of correlation. **C)** Difference in magnitude of correlation coefficients presented in panels A and B. **D)** Statistical comparison of correlations presented in panels **A** and **B**. Correlations were compared using the cocor package in R[5]. Boxes marked with an X denote insignificant differences between correlations after adjusting for multiple comparisons using the false discovery rate[6]. Significant differences in correlation strengths across datasets may be due to differences in dMRI acquisitions, differences in microstructural development, and other factors.

Abbreviations: ICP, inferior cerebellar peduncle; MCP, middle cerebellar peduncle; SCP, superior cerebellar peduncle; PF, parallel fibers; IP, input and Purkinje fibers; ABCD, Adolescent Brain and Cognitive Development Study; HCP-YA, Human Connectome Project-Young Adult dataset; FA, fractional anisotropy; MD, mean diffusivity; NoS, Number of Streamlines.

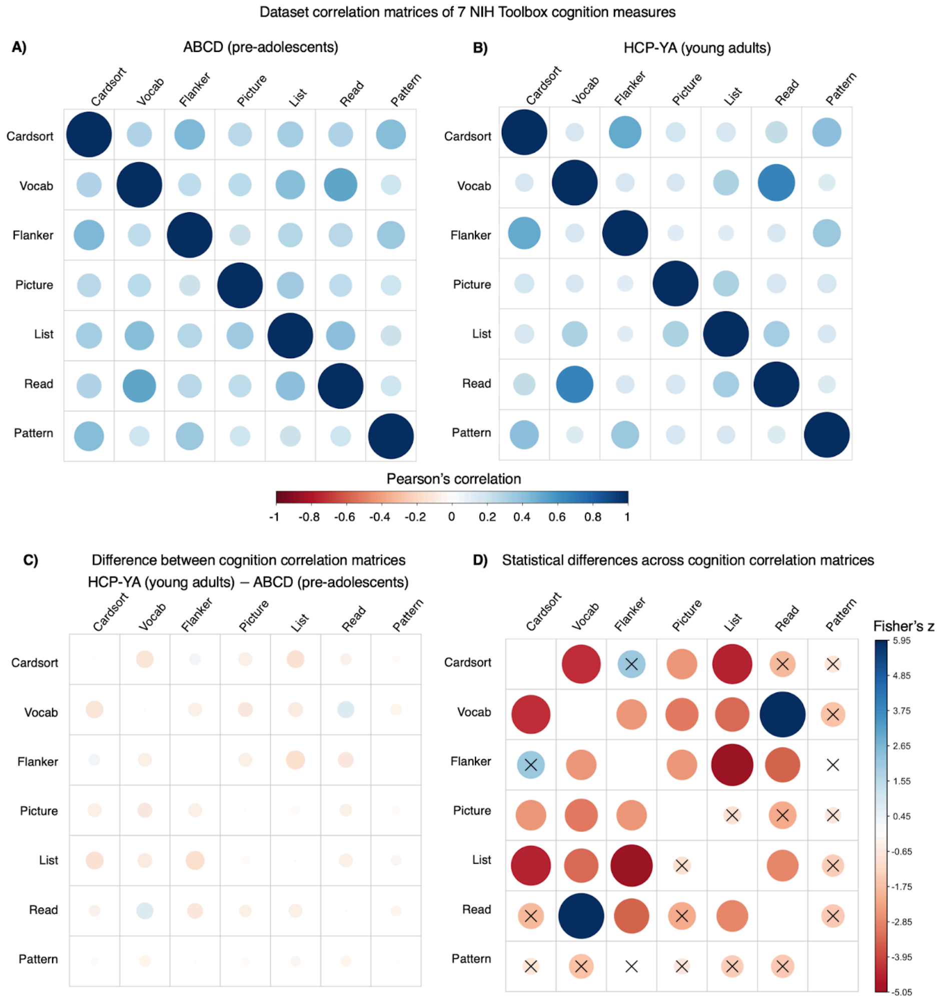


**SI Figure 4.** Dataset correlation matrices of cognition measures. Correlation matrices of seven NIH Toolbox cognition measures in **(A)** the ABCD dataset and **(B)** the HCP-YA dataset. Correlations were calculated using Pearson's correlation coefficient and plotted using the corrplot package in R [4]. Color indicates direction and magnitude of correlation. In both the ABCD and HCP-YA datasets, cognition measures are positively correlated across assessments. **C)** Difference in magnitude of correlation coefficients presented in panels **A** and **B**. **D)** Statistical comparison of correlations presented in panels **A** and **B**. Correlations were compared using the cocor package in R[5]. Boxes marked with an X denote insignificant differences between correlations after adjusting for multiple comparisons using the false discovery rate[6]. Significant differences in correlation strengths across datasets may be due to differences in data collection procedures (iPad vs. computer), differences in cognitive development (e.g., generalization vs. specialization), or other factors.

Abbreviations: HCP-YA, Human Connectome Project-Young Adult dataset; Vocab, NIH Toolbox Picture Vocabulary Test; Read, NIH Toolbox Oral Reading Recognition Test; Cardsort, NIH Toolbox Dimensional Change Card Sort Test; Flanker, NIH Toolbox Flanker Inhibitory Control and Attention Test; Pattern, NIH Toolbox Pattern Comparison Processing Speed Test; List, NIH Toolbox List Sorting Working Memory Test; Picture, NIH Toolbox Picture Sequence Memory Test.

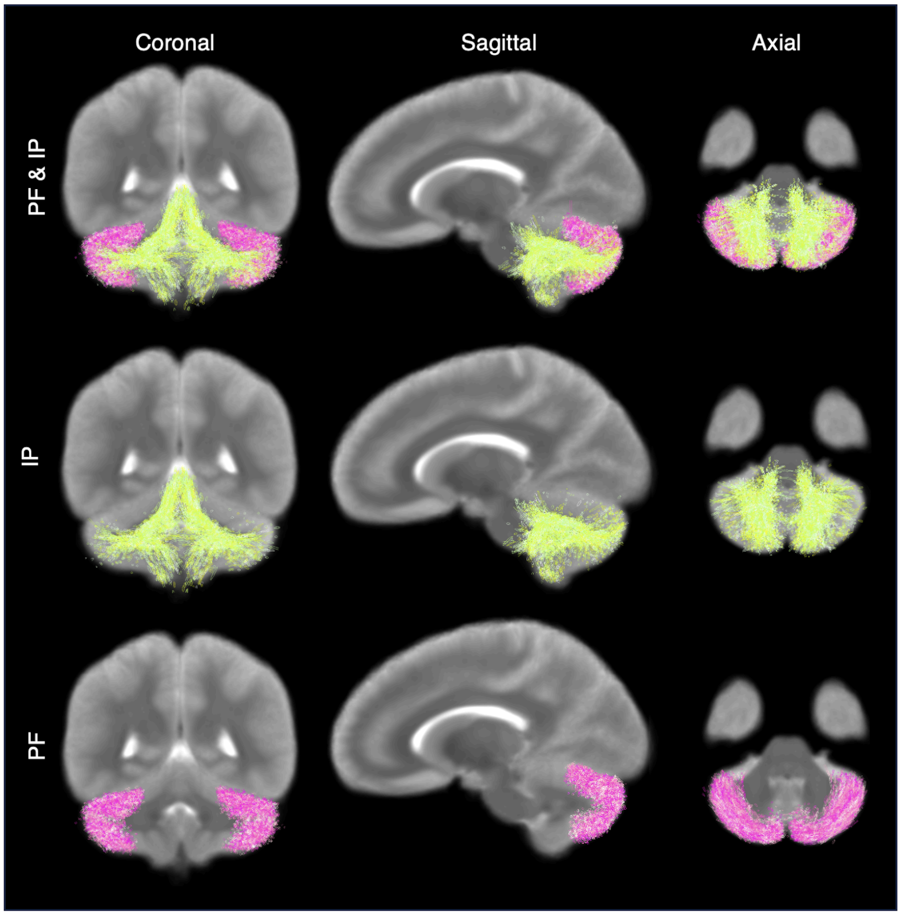


**SI Figure 5.** Anatomical location of the streamlines representing the parallel fibers (PF) and input and Purkinje fibers (IP). Coronal, sagittal, and axial views of the PF and IP as represented in the neuroanatomically curated pathway atlas. Fiber pathways are depicted in front of a T2 image slice. The PF is predominantly localized to the surface of the cerebellum, in the cortex. The IP fibers are located in the middle of the cerebellum, in the medullary white matter.

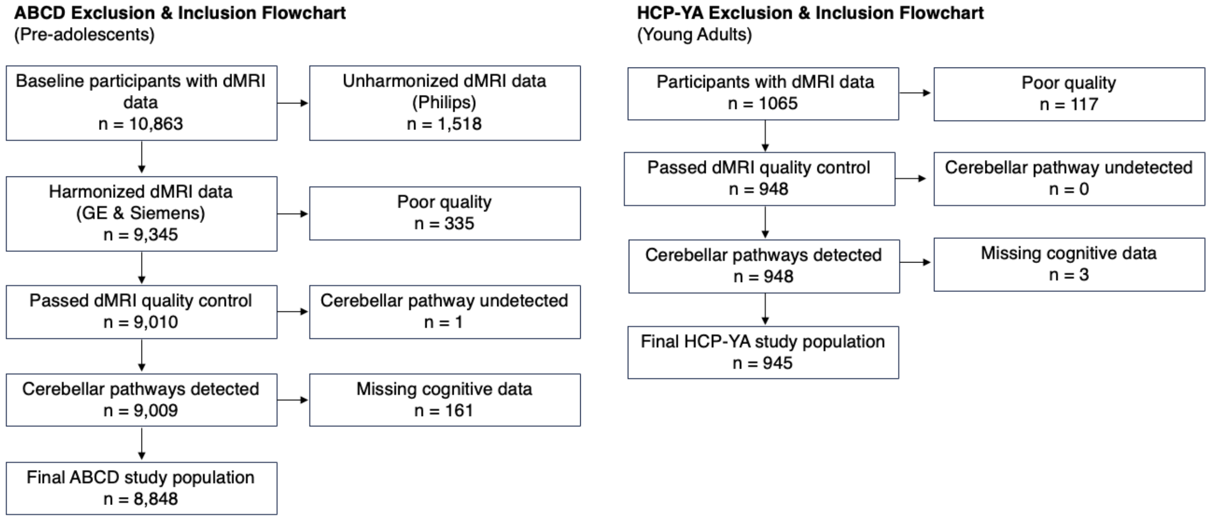


**SI Figure 6.** ABCD and HCP-YA study population exclusion and inclusion criteria.

### SI Tables

| dMRI Measure | Pathway | Mean | | Standard Deviation of Mean | |
|---|---|---|---|---|---|
| | | ABCD | HCP-YA | ABCD | HCP-YA |
| **FA** | ICP | 0.348 | 0.428 | 0.0263 | 0.0360 |
| | MCP | 0.558 | 0.580 | 0.0235 | 0.0301 |
| | SCP | 0.358 | 0.415 | 0.0219 | 0.0272 |
| | PF | 0.152 | 0.150 | 0.0122 | 0.0142 |
| | IP | 0.281 | 0.327 | 0.0180 | 0.0172 |
| **MD** | ICP | 0.000567 | 0.000508 | 0.0000181 | 0.0000124 |
| | MCP | 0.000551 | 0.000563 | 0.0000159 | 0.0000246 |
| | SCP | 0.000590 | 0.000559 | 0.0000148 | 0.0000157 |
| | PF | 0.000546 | 0.000514 | 0.0000134 | 0.0000149 |
| | IP | 0.000528 | 0.000473 | 0.0000130 | 0.0000113 |
| **NoS** | ICP | 651 | 1030 | 238 | 347 |
| | MCP | 1420 | 2800 | 430 | 910 |
| | SCP | 872 | 2200 | 219 | 420 |
| | PF | 6980 | 8120 | 3090 | 5170 |
| | IP | 2600 | 4630 | 708 | 1340 |

**SI Table 1.** Mean and standard deviation of dMRI measures of pathways in the ABCD and HCP-YA datasets.

Abbreviations: ICP, inferior cerebellar peduncle; MCP, middle cerebellar peduncle; SCP, superior cerebellar peduncle; PF, parallel fibers; IP, input and Purkinje fibers; ABCD, Adolescent Brain and Cognitive Development Study; HCP-YA, Human Connectome Project-Young Adult dataset; FA, fractional anisotropy; MD, mean diffusivity; NoS, Number of Streamlines.

| Cognitive Assessment | Mean | | Standard Deviation of Mean | |
|---|---|---|---|---|
| | ABCD | HCP-YA | ABCD | HCP-YA |
| Read | 90.95 | 116.90 | 6.81 | 10.50 |
| Vocab | 84.59 | 116.97 | 8.00 | 9.67 |
| Flanker | 94.27 | 111.66 | 8.95 | 10.20 |
| List | 96.96 | 111.22 | 11.80 | 11.40 |
| Cardsort | 92.72 | 115.07 | 9.37 | 10.20 |
| Pattern | 88.07 | 115.19 | 14.50 | 15.40 |
| Picture | 103.04 | 111.99 | 12.10 | 13.30 |

**SI Table 2.** Mean and standard deviation of unadjusted performance on NIH Toolbox cognitive assessments in the ABCD and HCP-YA datasets.

Abbreviations: ABCD, Adolescent Brain and Cognitive Development Study; HCP-YA, Human Connectome Project-Young Adult dataset; Vocab, NIH Toolbox Picture Vocabulary Test; Read, NIH Toolbox Oral Reading Recognition Test; Cardsort, NIH Toolbox Dimensional Change Card Sort Test; Flanker, NIH Toolbox Flanker Inhibitory Control and Attention Test; Pattern, NIH Toolbox Pattern Comparison Processing Speed Test; List, NIH Toolbox List Sorting Working Memory Test; Picture, NIH Toolbox Picture Sequence Memory Test.

| ABCD Dataset | | | | | |
|---|---|---|---|---|---|
| **NIH Toolbox Cognitive Assessment** | **Quantitative dMRI Measure** | **Cerebellar Pathway** | **Standardized *β*** | **95% Confidence Interval** | **p-value** |
| Cardsort | FA | IP | 0.03127 | [0.01021, 0.05232] | 0.008818** |
| Cardsort | FA | SCP | 0.07735 | [0.05616, 0.09854] | 9.433E-12*** |
| Cardsort | MD | SCP | 0.03619 | [0.01562, 0.05677] | 0.001654** |
| Cardsort | NoS | ICP | 0.06316 | [0.04277, 0.08355] | 1.064E-08*** |
| Cardsort | NoS | IP | 0.05552 | [0.03451, 0.07654] | 1.197E-06*** |
| Cardsort | NoS | SCP | 0.03107 | [0.01037, 0.05178] | 0.008173** |
| Flanker | FA | SCP | 0.05167 | [0.03035, 0.07299] | 9.006E-06*** |
| Flanker | MD | SCP | 0.03884 | [0.01818, 0.05951] | 0.0007128*** |
| Flanker | NoS | ICP | 0.05346 | [0.03297, 0.07396] | 1.544E-06*** |
| Flanker | NoS | IP | 0.04492 | [0.0238, 0.06604] | 0.0001047*** |
| List | FA | IP | 0.04159 | [0.02032, 0.06285] | 0.0004055*** |
| List | FA | SCP | 0.07779 | [0.05637, 0.0992] | 1.1E-11*** |
| List | MD | SCP | 0.04381 | [0.02302, 0.06459] | 0.0001194*** |
| List | NoS | ICP | 0.08626 | [0.06569, 0.1068] | 3.034E-15*** |
| List | NoS | IP | 0.06732 | [0.0461, 0.08854] | 4.588E-09*** |
| List | NoS | PF | 0.02813 | [0.00731, 0.04895] | 0.0189* |
| List | NoS | SCP | 0.03886 | [0.01794, 0.05977] | 0.0008171*** |
| Pattern | FA | SCP | 0.04746 | [0.02637, 0.06856] | 3.788E-05*** |
| Pattern | NoS | ICP | 0.04892 | [0.02863, 0.0692] | 9.716E-06*** |
| Pattern | NoS | IP | 0.02702 | [0.006104, 0.04793] | 0.02589* |
| Picture | FA | SCP | 0.05534 | [0.03385, 0.07683] | 2.082E-06*** |
| Picture | MD | SCP | 0.03363 | [0.01279, 0.05447] | 0.004326** |
| Picture | NoS | ICP | 0.0588 | [0.03815, 0.07946] | 1.531E-07*** |
| Picture | NoS | IP | 0.03151 | [0.01021, 0.05282] | 0.00895** |
| Vocab | FA | IP | 0.05635 | [0.03553, 0.07716] | 6.353E-07*** |
| Vocab | FA | SCP | 0.139 | [0.1182, 0.1599] | 1.031E-36*** |
| Vocab | MD | IP | 0.05378 | [0.03338, 0.07418] | 1.209E-06*** |
| Vocab | MD | PF | 0.05815 | [0.03784, 0.07845] | 1.333E-07*** |
| Vocab | MD | SCP | 0.09822 | [0.07794, 0.1185] | 7.185E-20*** |
| Vocab | NoS | ICP | 0.1013 | [0.08121, 0.1214] | 3.739E-21*** |
| Vocab | NoS | IP | 0.1002 | [0.07944, 0.1209] | 7.185E-20*** |
| Vocab | NoS | MCP | 0.04804 | [0.02759, 0.06849] | 1.695E-05*** |
| Vocab | NoS | PF | 0.04399 | [0.0236, 0.06437] | 8.265E-05*** |
| Vocab | NoS | SCP | 0.05563 | [0.03516, 0.0761] | 5.984E-07*** |
| Read | FA | ICP | 0.03507 | [0.01409, 0.05605] | 0.002994** |
| Read | FA | IP | 0.04751 | [0.02652, 0.06849] | 3.459E-05*** |
| Read | FA | MCP | 0.03199 | [0.0109, 0.05309] | 0.007584** |
| Read | FA | SCP | 0.1055 | [0.08438, 0.1265] | 4.819E-21*** |

| | | | | | |
|---|---|---|---|---|---|
| Read | MD | IP | 0.03231 | [0.01173, 0.05288] | 0.005622** |
| Read | MD | PF | 0.05941 | [0.03896, 0.07987] | 8.989E-08*** |
| Read | MD | SCP | 0.08856 | [0.06811, 0.109] | 3.666E-16*** |
| Read | NoS | ICP | 0.08937 | [0.06907, 0.1097] | 1.248E-16*** |
| Read | NoS | IP | 0.07985 | [0.05892, 0.1008] | 9.538E-13*** |
| Read | NoS | MCP | 0.06228 | [0.04169, 0.08287] | 2.376E-08*** |
| Read | NoS | PF | 0.03183 | [0.01128, 0.05238] | 0.006316** |
| Read | NoS | SCP | 0.04734 | [0.0267, 0.06798] | 2.721E-05*** |
| **HCP-YA Dataset** | | | | | |
| **NIH Toolbox Cognitive Assessment** | **Quantitative dMRI Measure** | **Cerebellar Pathway** | **Standardized *β*** | **95% Confidence Interval** | **p-value** |
| Cardsort | FA | SCP | 0.08374 | [0.01986, 0.1476] | 0.04394* |
| Cardsort | MD | ICP | 0.1006 | [0.03685, 0.1644] | 0.01245* |
| Cardsort | MD | PF | 0.09357 | [0.02863, 0.1585] | 0.02514* |
| Cardsort | MD | SCP | 0.0986 | [0.03559, 0.1616] | 0.01255* |
| Flanker | FA | PF | -0.1178 | [-0.1812, -0.05443] | 0.002095** |
| Flanker | MD | ICP | 0.1025 | [0.03926, 0.1658] | 0.009962** |
| Flanker | MD | PF | 0.1256 | [0.06143, 0.1898] | 0.00106** |
| Flanker | MD | SCP | 0.08678 | [0.0242, 0.1494] | 0.0316* |
| List | FA | SCP | 0.1254 | [0.0614, 0.1894] | 0.00106** |
| List | NoS | SCP | 0.1431 | [0.07746, 0.2087] | 0.0002717*** |
| Picture | FA | SCP | 0.09959 | [0.03573, 0.1634] | 0.01255* |
| Vocab | FA | SCP | 0.1967 | [0.134, 0.2594] | 1.009E-07*** |
| Vocab | MD | IP | 0.08945 | [0.02447, 0.1544] | 0.03208* |
| Vocab | MD | PF | 0.1362 | [0.07172, 0.2006] | 0.0003706*** |
| Vocab | MD | SCP | 0.1668 | [0.1046, 0.2291] | 3.143E-06*** |
| Vocab | NoS | IP | 0.08694 | [0.02043, 0.1535] | 0.04394* |
| Vocab | NoS | MCP | 0.132 | [0.06935, 0.1947] | 0.0003706*** |
| Vocab | NoS | SCP | 0.1991 | [0.1347, 0.2636] | 1.009E-07*** |
| Read | FA | MCP | 0.08932 | [0.02525, 0.1534] | 0.0316* |
| Read | FA | SCP | 0.171 | [0.1075, 0.2344] | 3.143E-06*** |
| Read | MD | IP | 0.1146 | [0.0493, 0.1799] | 0.004182** |
| Read | MD | PF | 0.1398 | [0.07487, 0.2047] | 0.0003041*** |
| Read | MD | SCP | 0.1745 | [0.1119, 0.2372] | 1.517E-06*** |
| Read | NoS | MCP | 0.1377 | [0.07463, 0.2008] | 0.0002717*** |
| Read | NoS | SCP | 0.1871 | [0.122, 0.2522] | 7.748E-07*** |

**SI Table 3.** Significant $\beta$ coefficients estimating relationships between quantitative dMRI measures of cerebellar pathways and NIH Toolbox cognitive assessments as measured in the ABCD and HCP-YA datasets. $\beta$ coefficients and p-values were calculated using multiple linear regression, controlling for age, sex (in the ABCD and HCP-YA datasets), and in-scanner head motion (ABCD dataset only). We report significant ($p< 0.05$) FDR corrected p-values; * < 0.05, ** < 0.01, *** < 0.001.

Abbreviations: ICP, inferior cerebellar peduncle; MCP, middle cerebellar peduncle; SCP, superior cerebellar peduncle; PF, parallel fibers; IP, input and Purkinje fibers; ABCD, Adolescent Brain and Cognitive Development Study; HCP-YA, Human Connectome Project-Young Adult dataset; FA, fractional anisotropy; MD, mean diffusivity; NoS, Number of Streamlines; Vocab, NIH Toolbox Picture Vocabulary Test; Read, NIH Toolbox Oral Reading Recognition Test; Cardsort, NIH Toolbox Dimensional Change Card Sort Test; Flanker, NIH Toolbox Flanker Inhibitory Control and Attention Test; Pattern, NIH Toolbox Pattern Comparison Processing Speed Test; List, NIH Toolbox List Sorting Working Memory Test; Picture, NIH Toolbox Picture Sequence Memory Test.

| **Cerebellar Pathway** | **Atlas Fiber Clusters** |
|---|---|
| SCP | cluster_00118, cluster_00120, cluster_00493, cluster_00566 |
| MCP | cluster_00110, cluster_00111, cluster_00115,cluster_00520, cluster_00523, cluster_00526, cluster_00544, cluster_00546, cluster_00550 |
| ICP | cluster_00126, cluster_00130, cluster_00515 |
| PF | cluster_00495, cluster_00500, cluster_00502, cluster_00503, cluster_00505, cluster_00508, cluster_00510, cluster_00512, cluster_00513, cluster_00516, cluster_00519 cluster_00521, cluster_00528, cluster_00529, cluster_00536, cluster_00538, cluster_00539, cluster_00541, cluster_00548, cluster_00551, cluster_00552, cluster_00562, cluster_00563, cluster_00565, cluster_00571 |
| IP | cluster_00109, cluster_00496, cluster_00498, cluster_00501, cluster_00504, cluster_00514, cluster_00527, cluster_00537, cluster_00540, cluster_00570, cluster_00572, cluster_00575 |

**SI Table 4.** Fiber clusters from the O'Donnell Research Group (ORG) Fiber Clustering White Matter Atlas that form the investigated neuroanatomical cerebellar pathways. We used the ORG-800FC-100HCP version of the atlas which is available online (https://github.com/SlicerDMRI/ORG-Atlases)[7].
